# Spacing of Cue-approach Training Leads to Better Maintenance of Behavioral Change

**DOI:** 10.1101/268821

**Authors:** Akram Bakkour, Rotem Botvinik-Nezer, Neta Cohen, Ashleigh M. Hover, Russell A. Poldrack, Tom Schonberg

**Author notes:** Current Address: Department of Psychology, Columbia University, New York, NY, USA. Current Address: Department of Psychology, Stanford University, Stanford, CA, USA. Corresponding authors: (TS), (AB).

## Abstract

The maintenance of behavioral change over the long term is essential to achieve public health goals such as combatting obesity and drug use. Previous work by our group has demonstrated a reliable shift in preferences for appetitive foods following a novel non-reinforced training paradigm. In the current studies, we tested whether distributing training trials over two consecutive days would affect preferences immediately after training as well as over time at a one-month follow-up. In four studies, three different designs and an additional pre-registered replication of one sample, we found that spacing of cue-approach training induced a shift in food choice preferences over one month. The spacing and massing schedule employed governed the long-term changes in choice behavior. Applying spacing strategies to training paradigms that target automatic processes could prove a useful tool for the long-term maintenance of health improvement goals with the development of real-world behavioral change paradigms that incorporate distributed practice principles.

## Introduction

The potential for targeting automatic processes to change human behavior has become increasingly clear [1], especially in light of the relative ineffectiveness of relying on effortful control of behavior, given the largely automatic and habitual nature of everyday human behavior [2]. Previous research aimed at changing choice preferences for appetitive foods employed a novel non-reinforced training paradigm named “cue-approach training” [CAT, 3]. Cue-approach training was found to be effective at influencing choice behavior in an immediate test [3–7] and the choice preference shift was shown to persist over at least two months after the longest cue-approach training employed [3,8]. The cue-approach task is related to previous work showing that visual attention both reflects and influences choices [9–11] and to other research on the attentional boost effect that highlights the importance of driving attention at behaviorally relevant points in time in boosting memory for incidental stimuli [12–14]. During CAT, participants learn to associate a tone cue to press a button when particular food items appear on the screen. We have posited that heightened attention during behaviorally relevant points in time modulates the subjective value placed on cue-associated foods [3,4]. Although the specific contribution of memory to the cue-approach choice effect remains unknown, the non-externally reinforced associative nature of cue-approach training suggests that memory principles apply to the cue-approach task [5]. The specific cognitive and neural mechanisms by which CAT leads to a change in choice preferences is under active investigation. We previously proposed and provided evidence to support our hypothesis that CAT drives attention toward particular stimuli when cued to make an approach response. The task potentially also engages cognitive control mechanisms that interact with valuation mechanisms and modulate the value of those same items, which manifests as a shift in choice preferences [3-5]. It remains possible - indeed likely - that the stimulus-cue-response association formed during CAT is retrieved when a stimulus appears paired with another non-cued item during the choice phase to bias preference to the cued item. CAT can be described as the mirror task to the automated inhibition version of the stop-signal task (SST) [15]. The SST is an associative learning task that has been shown to slow reaction times to stimuli previously associated with inhibition of a pre-potent response, suggesting retrieval of stimulus-stop associations during the test phase. We have previously employed a training phase identical to SST in a cue-avoidance (rather than cue-approach) design to test for avoidance-tendencies during choice, but found no effect, suggesting that approach-, rather than avoidance-tendencies are modulated during CAT [3]. Given the potential importance of the stimulus-cue-response association in CAT, we hypothesized that distributed training - or spacing of training trials - will lead to a more robust and lasting shift in choice preferences. In the studies described here, we distributed CAT training trials over two days to test the effects of this training schedule on maintenance of the preference shift one week and one month after initial training.

One of the oldest and most reliable findings in research on human learning is the spacing effect. Ebbinghaus [16] was the first to report, over 100 years ago, the benefits of spacing trial presentations in time during learning on subsequent retrieval strength. Hundreds of studies have since established the spacing effect as a potent tool to improve memory retention [for review see 17,18]. Information studied across multiple sessions, spaced out over days or weeks, is often better retained than information studied in the same amount of time in a single session when tested for weeks or months after the end of the study phase. When Reynolds and Glaser [19] taught unfamiliar biology terms to student participants over multiple consecutive repetitions in a single 40 minute classroom science period (massed learning) or spaced the review over multiple classroom science periods with other learning tasks in the intervening class periods (spaced learning), they found that spacing review over time produced significantly better retention of the material. Meta-analysis of the spacing effect in verbal learning tasks revealed that spaced (vs. massed) learning of items consistently leads to better long-term retention [18]. Spacing repetitions every few minutes has the most benefit for retention on a test one day later whereas spacing repetitions every day has the most benefit for retention over days or weeks [18].

Researchers have also manipulated the lag, or length of spacing between study presentations. The lag effect refers to improvements in memory performance for information that was repeated over longer lags compared to information repeated over shorter lags. Madigan [20] gave participants lists of words, some of which were presented twice. The lag, or number of intervening words between the two presentations, varied. Recall for repeated words improved with longer lags. Here, for the sake of simplicity, we refer more generally to the spacing effect as the benefit of longer spacing on memory retention, where spaced training trials are distributed over longer time periods (in our case two consecutive days) versus massed trials that are distributed over shorter time periods (in our case on a single day).

The spacing effect has been demonstrated across a number of types of learning. In addition to the meta-analysis of the spacing effect on verbal learning conducted by Cepeda et al. [18], Lee and Genovese [21] conducted a meta-analysis examining the effects of spacing practice on motor skills. They found that spaced practice enhances acquisition of motor skills compared to massed practice but more importantly it resulted in greater retention of motor skills compared to massed practice. Spacing strategies have also been successfully implemented to reduce the return of fear in treatment of anxiety disorders [22]. Participants with public speaking anxiety who underwent a spaced schedule of exposure therapy experienced less return of fear at one-month follow-up than matched participants who followed a massed therapy schedule.

Based on all of this research, spacing treatment sessions holds great promise to help maintain behavioral change over longer terms than massed training. To our knowledge, this strategy has not yet been applied to other behavioral change efforts outside the fear domain. The goal of the current studies was to test whether spaced training of CAT over two days results in longer retention of a shift in choice preferences after one week and one month compared to when the training was massed in a single training session. We hypothesized that spaced training would lead to better maintenance of the observed shift in choice preferences. To test this, individual items during the training phase were either Go (paired with the cue to respond) or NoGo (not cued, two Go factor levels) and either spaced (trained over two days) or massed (trained on a single day, two spacing factor levels). The probe choice phase pitted two foods against each other and was repeated immediately after training, one week, and one month later (three time factor levels). Given limitations on the number of pairs of items we are able to form and to optimize power to detect an effect, we did not test a full 2 × 2 × 3 design. Instead, we tested two separate 2 × 3 designs. The first was 2 spacing × 3 time points. We kept spacing status constant across the two items in a pair and items differed only on Go status. We hypothesized that choices for Go over NoGo items would remain higher over time when both items were Spaced compared to when both items were Massed. More specifically, we hypothesized an interaction between spacing and time factors on the choice of Go over NoGo driven by a constant rate of Go choices over time when both items are spaced and a decreasing rate of Go choices over time when both items are massed. If there is no interaction, we hypothesized that there would be a main effect of Spaced greater than Massed for choices of Go over NoGo. As for the 2 Go status × 3 time points design, both foods were Go or NoGo but differed on training spacing schedule. Here, we hypothesized that choices for Spaced over Massed items would remain higher or perhaps increase over time when both items are Go compared to when both items were NoGo. Specifically, we hypothesized an interaction between Go and time factors on the choice of Spaced over Massed driven by an increasing rate of choices for Spaced over time when both items are Go and a constant rate when both items are NoGo. If there is no interaction, we hypothesized that there would be a main effect of Go greater than NoGo for choices of Spaced over massed. The studies presented here apply the principles of spaced learning to a non-reinforced training task that targets automatic cognitive processes (CAT) and has been proven to influence appetitive choice behavior [3–7] to test the effectiveness of distributed practice on the maintenance of behavioral change over time.

## Materials and Methods

### Overview

In the studies reported here, we spaced cue-approach training over two consecutive days to test whether spacing improves the maintenance of a shift in choice behavior. In the standard cue-approach task, single images of food are presented one at a time and participants are instructed to press a button on the keyboard as fast as they can when they hear a neutral tone. The cue tone is paired with some items (Go items) and not with others (NoGo items). In a subsequent choice phase, participants choose between two items. Different pairs of items appear on each trial, but each pair contains one Go and one NoGo item that were equated for pre-experimental preferences. Participants are told that they will receive the item they chose on a randomly selected trial. In this phase, participants reliably tend to choose Go over NoGo items [3].

To facilitate discussion of methods and results across the four studies presented here, we define a Spaced item as an item that appeared on both days of cue-approach training (i.e. half of the training phase presentations were on day 1 and the second half of the training phase presentations appeared on day 2). We define Massed items as items that were trained on a single day, i.e. all the training phase presentations appeared on the same day. We define within-session lag as the average number of intervening other-item trials between presentations of a particular item on one day.

In the four studies reported here, we tested the effect that spacing cue-approach training over two consecutive days had on choice behavior after one week and one month. Additionally, we tested the effect of the order in which Massed items appeared: either on the first or second training day. Finally, we tested how the length of within-session lag during training influenced the choice effect.

### Participants

A total of 114 healthy young participants took part in one of four spaced CAT studies. Table 1 summarizes participant demographic characteristics for the four studies. Sample sizes are similar to previous studies and were determined prior to data collection. No statistical methods were employed to determine the sample size for the first three studies. Power analysis was used to determine the sample size for the fourth study (replication of study 3) based on effect sizes from study 3. The power analysis determined that 35 participants were sufficient to detect the effects of interest from study 3. We over-recruited to anticipate attrition over the four study sessions, but we overestimated the rate of attrition and are including 39 participants that completed study 4. Results remain the same if only the initial 35 participants are included.

**Table 1:**
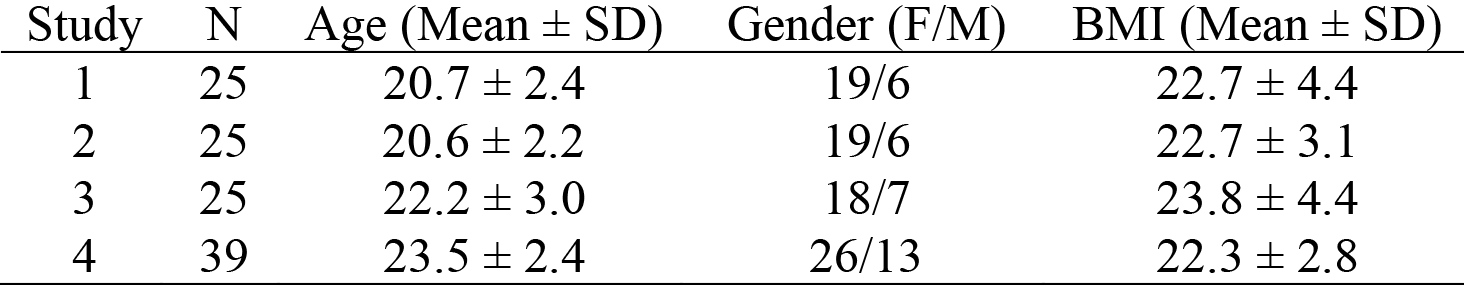
Demographic details for all studies. SD (Standard Deviation). BMI (Body Mass Index).

All participants fit the same inclusion criteria as previously described [4] (i.e., they had normal or corrected-to-normal vision, no history of psychiatric, neurologic or metabolic illness, no history of eating disorders, no food restrictions, and were not taking any medication that would interfere with the experiment). During recruitment, participants were told that our goal was to study food preferences and were asked to fast for four hours prior to each of their visits to the laboratory. Participants were also informed of and scheduled for four visits to the lab: two initial visits on two consective days, a follow-up visit one week after day 2 (day 9), and a final visit four weeks after day 2 (day 30). The study was approved by the institutional review board (IRB) at the University of Texas at Austin and at Tel Aviv University (study 4), and all participants gave informed consent.

### Stimuli

Color photographs of 60 appetitive junk food items were used. These stimuli were used in previous studies [3,23]. The snacks for sample 4 were of Israeli snacks and can be found in schonberglab.tau.ac.il/resources and were previously used [8].

## General procedure across studies

### Auction

The auction procedure (Fig 1A) was described in detail in previous publications [3–5] and followed the procedure of a Becker-Degroot-Marschak (BDM) auction [24]. Briefly, single pictures of food items were presented on the screen and participants indicated their willingness-to-pay (WTP) for each individual item by selecting a value on a visual analog scale placed at the bottom of the screen using the computer mouse. The experimenter explicitly explained to participants that the best strategy was to bid exactly what the item was worth to them to buy from the experimenter at the end of the session. At the end of the session, the computer generated a counter bid (a random number between 0 and 3 in 25 cent increments) and compared it to the participant’s bid on a randomly drawn auction trial. If the participant’s bid was lower than the computer’s, they lost the auction and could not buy that item. If the participant bid the same or outbid the computer, then they were offered that item at the computer’s bid lower price. We then used WTPs to rank order all 60 foods for each participant from most preferred (highest WTP) to least preferred (lowest WTP, Fig 2A). Items were split into higher-value and lower-value items according to the median. Items were then assigned to one of two training conditions; Go items required a button press during training and NoGo items required no response from the participant in adaptations of the cue-approach task [3]. Item assignment to Go and NoGo conditions based on their rank order was counterbalanced across participants.

**Fig 1:**
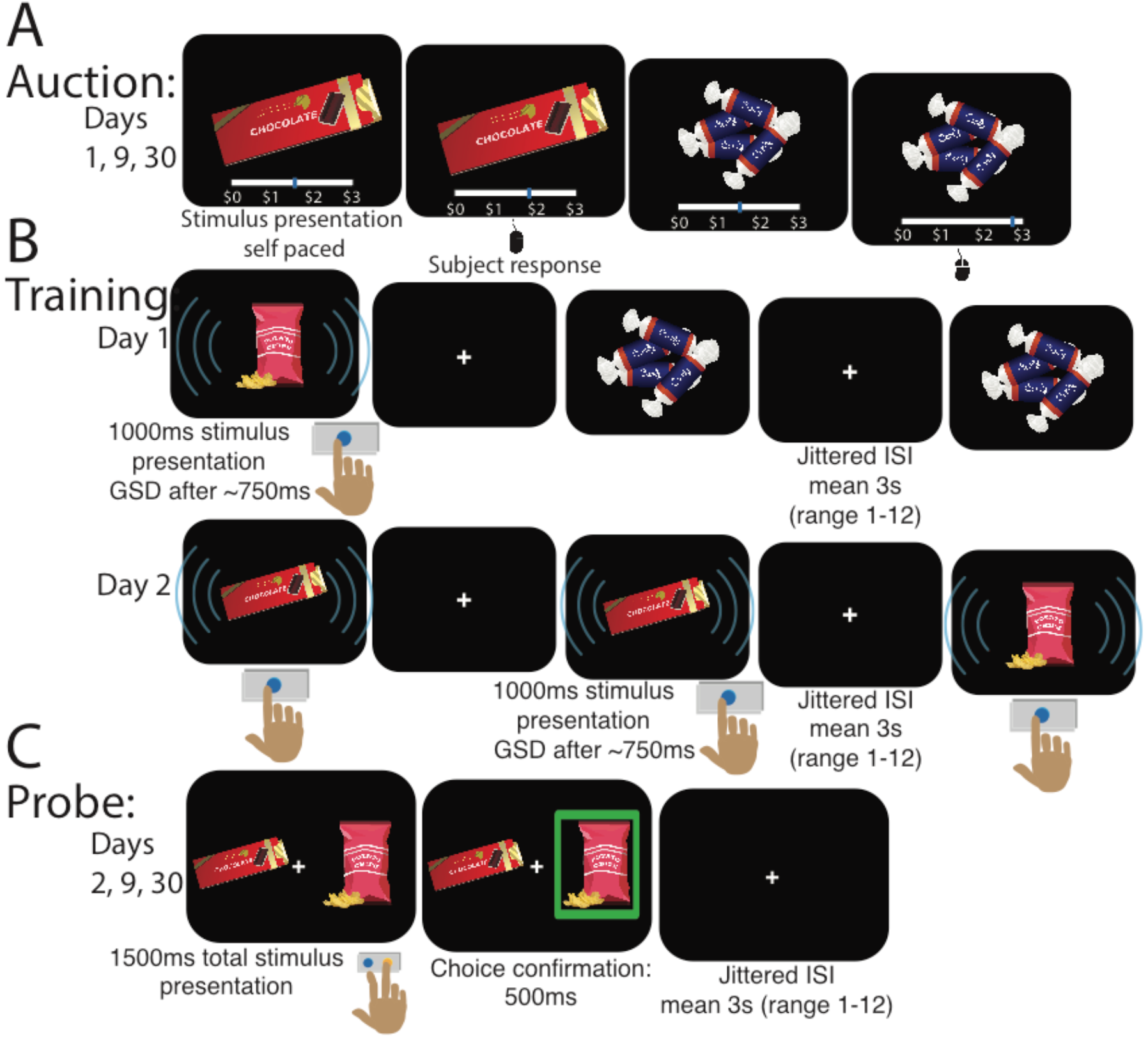
Spacing of cue-approach training trials task procedure. A) Participants first take part in an auction. Food items are then rank ordered based on WTP and assigned to one of four conditions (Fig 2). B) Participants are then asked to observe the items and to press a button as quickly as possible only when they hear an infrequent tone (GO items). The tone sounds at a variable time after the food stimulus appears on the screen (GO signal delay [GSD]). GSDs are adjusted using a staircase procedure. Spaced items are seen on both day 1 and day 2 of training (e.g. potato chips). Massed items are trained only on one of the days (e.g. candy trained only on day 1 or chocolate trained only on day 2). C) Participants then choose between two items that are matched for WTP but differ on Go/NoGo or Spaced/Massed status for consumption in a probe phase. Participants then return for two follow-up probes one week and one month later.

**Fig 2:**
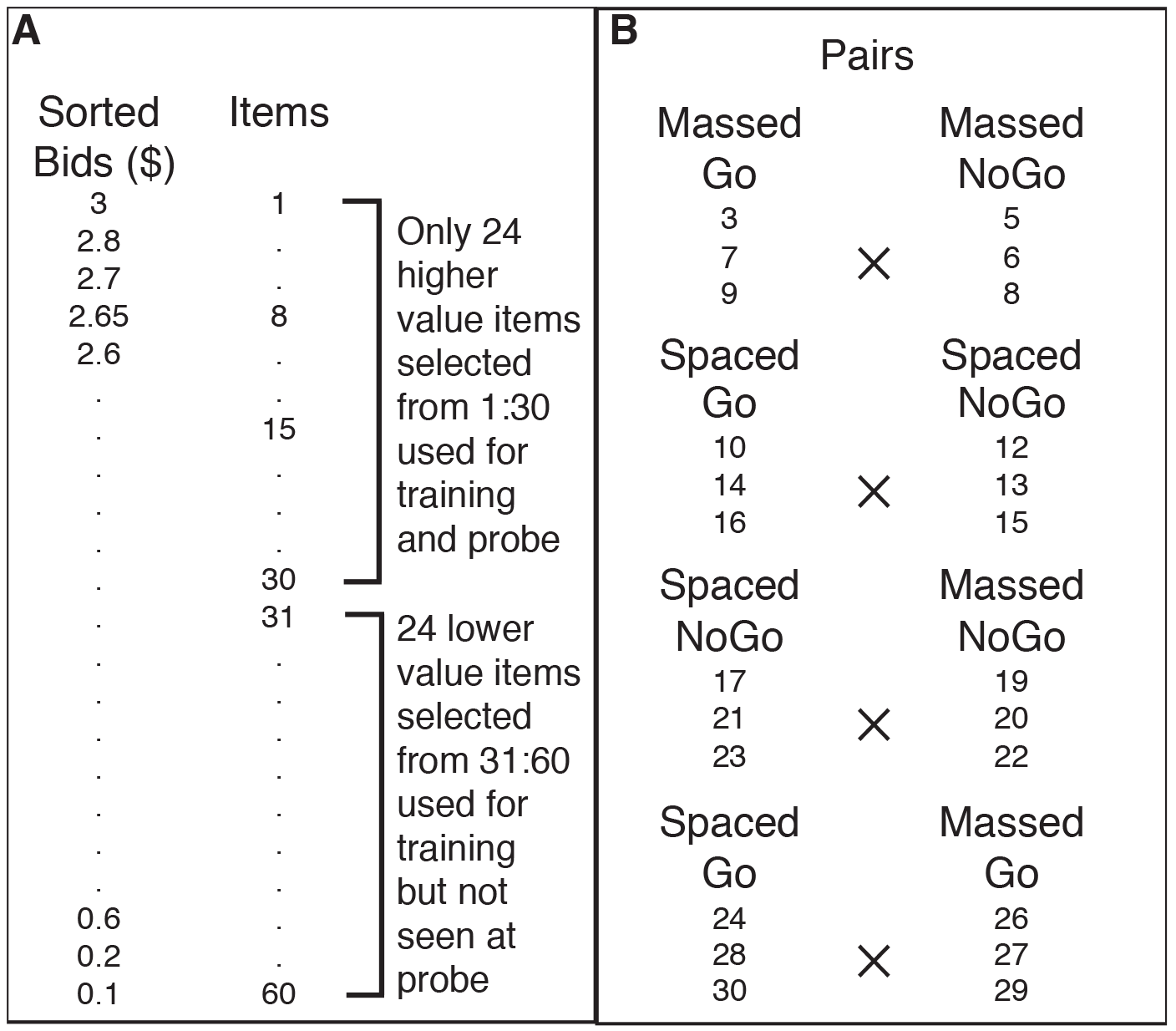
Sorting and pair matching procedure used for all experiments. A) Items are rank ordered based on bid obtained in the first auction. Only 24 higher valued items (rank orders 3 to 30) are selected for use in both the training and probe phases (rank orders 4, 11, 18 and 25 not used). 24 lower valued items (from rank orders 31 to 60) are randomly selected and used as NoGo items during training but are not seen during probe. B) Probe items are assigned to one of four training conditions in a 2×2 design (Spaced Go, Massed Go, Spaced NoGo, Massed NoGo). Item condition assignments are counterbalanced across participants. Probe items are paired such that each pair contains two items that are matched for pre-experimental preference based on the initial auction to allow for four types of comparisons (Massed Go vs. Massed NoGo, Spaced Go vs. Spaced NoGo, Spaced NoGo vs. Massed NoGo and Spaced Go vs. Massed Go).

### Item selection

All 60 food items were rank ordered based on WTP from highest to lowest (Fig 2A). Twenty four of the 30 high-value items were placed in one of four training conditions in a 2 × 2 design (Go/NoGo × Spaced/Massed, Fig 2B) to allow for four different comparisons at probe: 1) Massed Go vs. Massed NoGo, 2) Spaced Go vs. Spaced NoGo, 3) Spaced NoGo vs. Massed NoGo, 4) Spaced Go vs. Massed Go. Each pair type at probe was composed of nine unique pairs. This item selection procedure ensured that items that were paired during probe were equated for WTP such that participants should be indifferent to the choice between items in each pair given their stated pre-experimental preferences. In order to ensure that only 25% of all trials were Go trials, in accordance with standard cue-approach training, we included 24 randomly selected low-value items (from the bottom 30 items in Fig 2A) as Spaced and Massed NoGo items during training, but these items (along with the 6 non-selected high-value items) were never seen during probe. Item assignment (based on rank order number) to each of the four training conditions (Go/NoGo × Spaced/Massed) for items that appeared during probe was counterbalanced across participants.

### Training

After completing the auction, participants started cue-approach training. They were asked to press a button on the keyboard as quickly as possible when they heard an infrequent neutral tone (Fig 1B). The general cue-approach training procedure is described in detail in Schonberg et al.[3]. The tone appeared at the beginning of training 750 ms after the food stimulus appeared on the screen and this Go signal delay (GSD) was adjusted using a staircase procedure to ensure that the participants would only achieve roughly 75% Go success, i.e. pressing the button after the tone sounds, but before the food stimulus disappears from the screen a fixed one second after the onset of a food stimulus. In all studies 12 out of a total of 48 trained items were consistently associated with a tone (Go items) during the training phase, which spanned two consecutive days. Items were either trained all on the same day (Massed items) or had half their training presentations on day 1 and the rest of the training presentations on day 2 (Spaced items). Spaced and Massed item presentations as well as Go and NoGo item presentations were intermixed. All items were presented a total of 12 times each during training. The three studies presented here differ on the days on which Massed items are trained (Table 2). These studies were designed to test potential primacy and recency effects as well as lag effects on choice during probe following spaced cue-approach training.

**Table 2:** Number of items (Items), number of presentations of each item (Pres.) and average within-session lag (Lag) for Spaced and Massed items on Day 1 and Day 2 of CAT (Fig 2B) for all studies.

| Study | Day 1 |  |  |  |  |  | Day 2 |  |  |  |  |  |
| --- | --- | --- | --- | --- | --- | --- | --- | --- | --- | --- | --- | --- |
|  | Spaced |  |  | Massed |  |  | Spaced |  |  | Massed |  |  |
|  | Items | Pres. | Lag | Items | Pres. | Lag | Items | Pres. | Lag | Items | Pres. | Lag |
| 1 | 24 | 6 | 24 | 0 | 0 | 0 | 24 | 6 | 72 | 24 | 12 | 36 |
| 2 | 24 | 6 | 72 | 24 | 12 | 36 | 24 | 6 | 24 | 0 | 0 | 0 |
| 3 & 4 | 24 | 6 | 48 | 12 | 12 | 0 | 24 | 6 | 48 | 12 | 12 | 0 |

### Probe

After filling out a computer adapted version of the Barratt Impulsiveness Scale [BIS-11, 25], or ranking fractal art images in study 4, participants were presented with pairs of items that were matched for WTP (based on the initial auction, Fig 2) and they were asked to choose one on each trial (Fig 1C). They were told that a single trial would be selected and honored for real at the end of the session, meaning they would receive that item to eat. Four types of pairs were formed based on training (Fig 2B). Each pair type was made up of nine unique pairs. Each unique pair was formed from three items, each paired with three other items with similar WTP but differed on one of the two factors: spacing or tone-pairing (see Fig 2B).

One week later, participants returned for a third visit (Fig 3). They performed another probe phase with the same pairs as in the first session but in a randomized trial order. They then took part in another auction, identical to the first, but with randomized trial order (Fig 3). The second auction allowed us to examine changes in WTP due to CAT. Finally, participants completed a memory task that assessed whether they could remember whether an item had been associated with a tone during training (was a Go item) or not (was a NoGo items).

**Fig 3:**
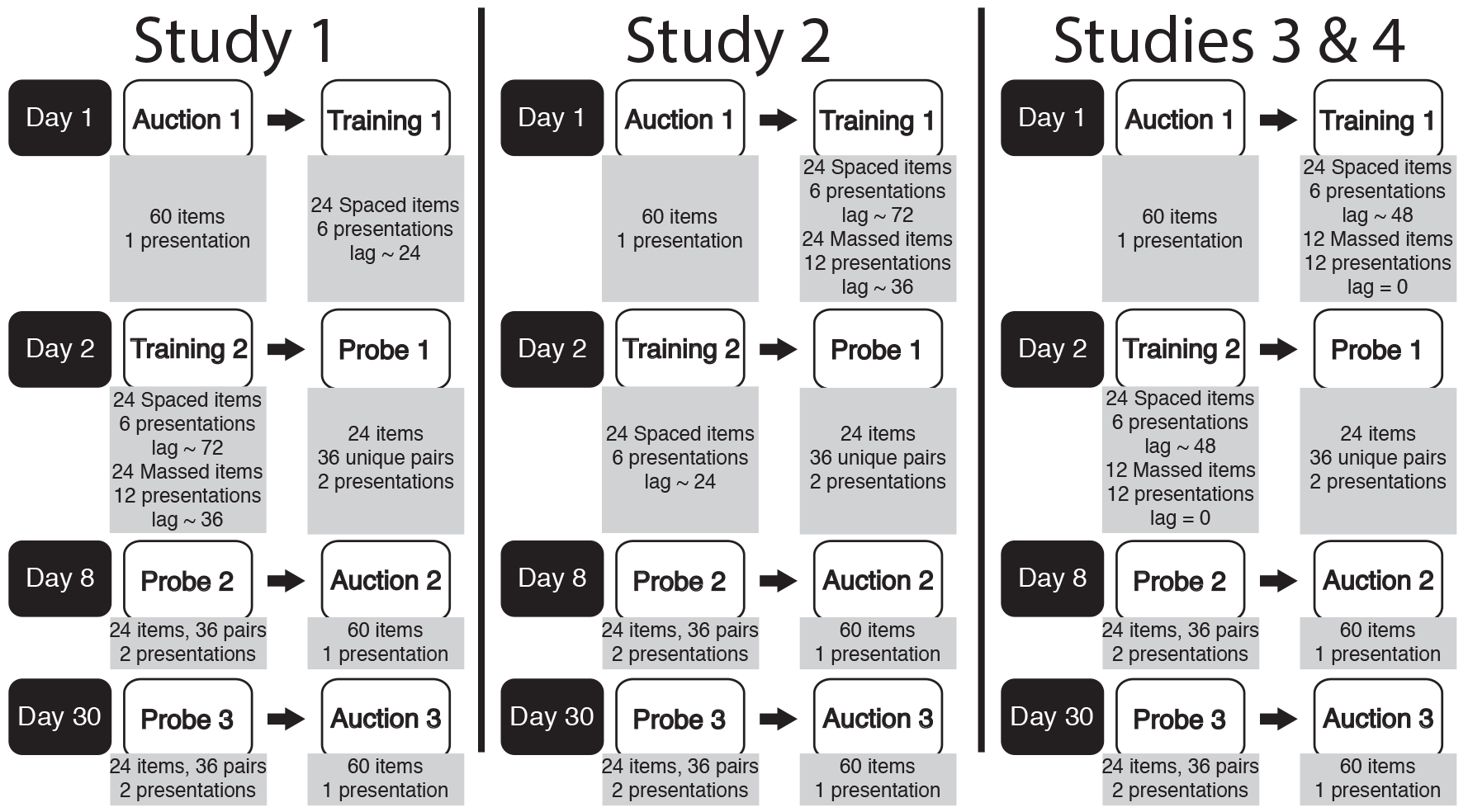
Spacing of cue-approach study layout over 30 days. On Day 1, participants take part in an initial auction (Fig 1A). They then start training on 24, 48 or 36 items (in Study 1, 2 and 3 & 4, respectively, Fig 1B, Table 2). On Day 2, participants return to continue training, now on 48, 24 and 36 items (in Study 1,2 and 3 & 4, respectively, Fig 1B, Table 2). They then make food choices in a first probe phase (Fig 1C). 24 items in 36 unique pairs are presented during probe. Participants return one week later on Day 9 and one month later on Day 30 and repeat the probe, identical to the first probe, but with random trial order. Finally, participants take part in an auction identical to the first, but with random trial order.

Approximately one month after the first visit, participants returned to the lab for a fourth visit (Fig 3). This visit was structured the same as the third visit. Visits three and four allowed us to examine the effectiveness of spaced CAT on the maintenance of choice preference for Go items and any induced choice preference for Spaced items.

## Specific methods for each study

### Study 1

During cue-approach training, only half the training presentations of all Spaced items were presented during day 1. The second half of the training presentations of all Spaced items appeared on day 2. All training presentations of Massed items appeared only on day 2. Because a particular massed item was presented twice for every presentation of a particular spaced item during training on day 2, within-session lag for Massed items - i.e. the average number of intervening trials between presentations of the same item - was shorter (~36 trials between same item presentations) than for Spaced items (~72 trials). Because Spaced items were presented both on day 1 and day 2, whereas Massed items only appeared on day 2, within-session lag for Spaced items expanded from day 1 (~24 trials) to day 2 (~72 trials between same item presentations). Auction and probe procedures remained the same across studies.

### Study 2

Participants in study 2 were trained on all Massed items only on day 1 of training, i.e. all training presentations of all Massed items took place on day 1. Only half the training presentations of all Spaced items took place on day 1. The second half of the training presentations of all Spaced items was presented during day 2. Because Massed items were presented twice for every presentation of a Spaced item on day 1 and Spaced items were presented on both days, within-session lag for Massed items was shorter (~36 trials between same item presentations) than for Spaced items (~72 trials). Because Massed items were only presented on day 1, within-session lag for Spaced items contracted from day 1 (~72 trials) to day 2 (~24 trials between same item presentations). Auction and probe procedures are described in the General Methods section above and remained the same across all studies.

### Studies 3 & 4

Participants in studies 3 & 4 had all 12 presentations of half the Massed items appear on day 1 of training and all 12 presentations of the second half of Massed items appear on day 2. Half of the presentations of all Spaced items were presented during day 1 and the second half of presentations of all Spaced items were presented during day 2 (Fig 3, Table 2). Massed items were zero-lag massed with zero within-session lag (i.e. presentations of a particular Massed item were presented consecutively with no other intervening items). Within-session lag for a particular Spaced item averaged 48 other item presentations and remained the same from day 1 to day 2.

## Analysis

### Probe

We predicted that distributing CAT over two consecutive days would induce lasting preference change in the form of greater choice of Spaced Go items. To test our prediction, we performed repeated measures logistic regression in two 2 × 3 designs. The first analysis tested the main effects of spacing (Spaced versus Massed) and time of test since training (linear effect from immediate to 1 week and 1 month) as well as the interaction of spacing and time for choice trials that pitted two foods that differed on training Go status (testing odds of choosing Go over NoGo against equal odds) but both items were trained using the same spacing schedule (both items were Spaced or both items were Massed, panels A in Figs 4–7). We also tested the simple effects and ran the regression separately for each pair type (Massed Go versus Massed NoGo & Spaced Go versus Spaced NoGo) for each probe (immediate, one week and one month follow-ups). The second analysis tested the main effects of Go status (Go vs. NoGo) and time of test since training (linear effect from immediate to 1 week and 1 month) as well as the interaction of spacing and time for choice trials that pitted two foods that differed on spacing schedule (testing odds of choosing Spaced over Massed against equal odds) but both items had the same Go status (both items were Go or both items were NoGo, panels B in Figs 4–7). We also tested the simple effects and ran the regression separately for each pair type (Spaced NoGo vs. Massed NoGo & Spaced Go vs. Massed Go) for each probe (immediate, one week and one month follow-ups). The experiment was not set up as a 2 Go status × 2 Spacing status × 3 Time design. By experimental design, the participants never encountered a Spaced Go with a Massed NoGo or a Massed Go with a Spaced NoGo in a pair on any trial.

**Fig 4:**
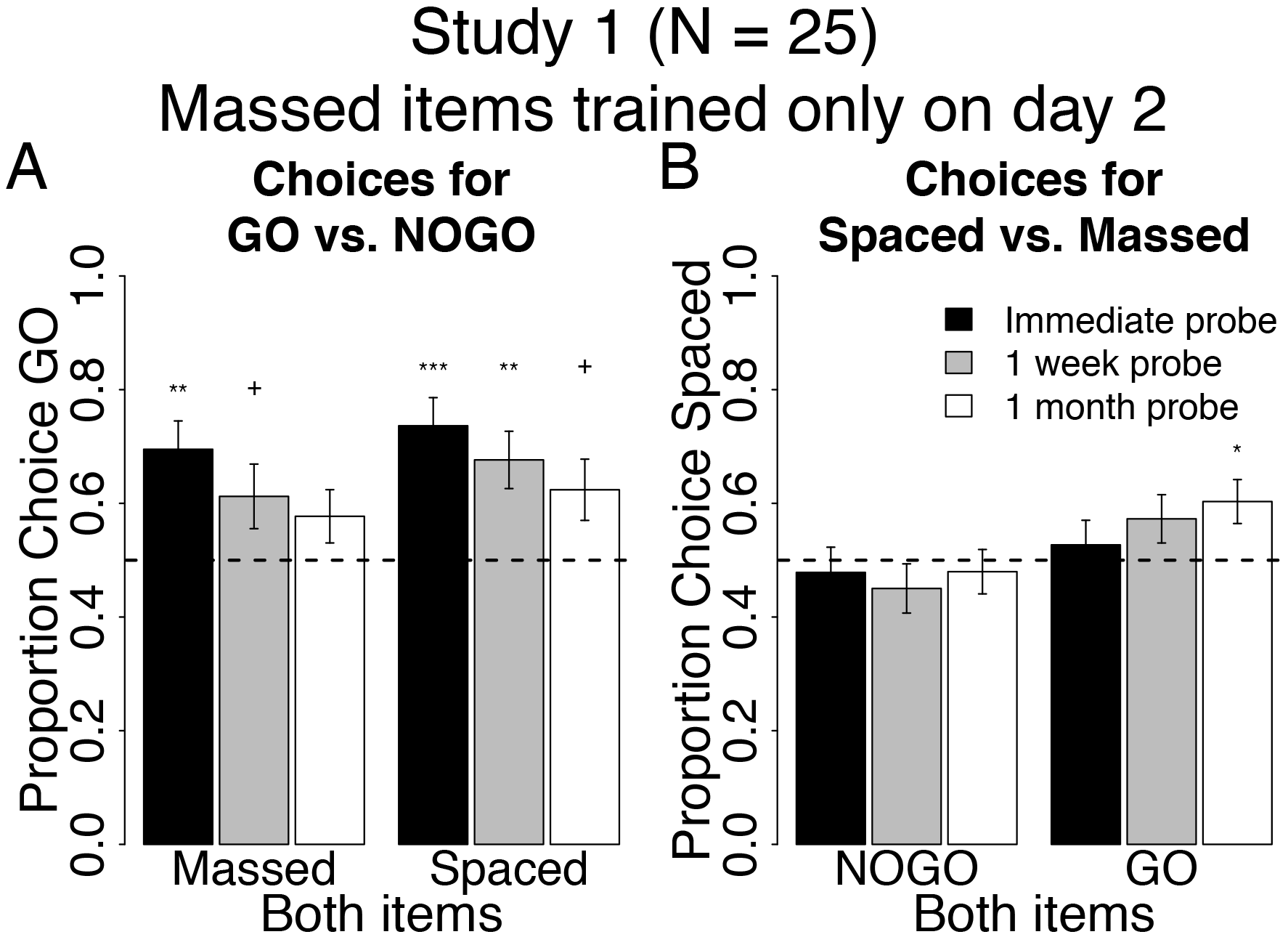
Behavioral results at probe for spacing of cue-approach training Study 1. Proportion choice of the Go vs. NoGo item (A) and proportion choice of Spaced vs. Massed item (B) at probe immediately after training (black bars), one week (grey bars) and one month later (white bars). All error bars reflect one standard error of the mean (SEM). +: *p* < 0.05, *: *p* < 0.01, **: *p* < 0.001, ***: *p* < 0.0001 in two-tailed repeated measures logistic regression for odds of choosing Go to NoGo (A) or Spaced to Massed (B) against equal odds.

### Auction

To test any change in the subjective value placed on individual items due to spaced cue-approach training, we used repeated-measures linear regression to test the two-way interaction between time (first, second [at one week follow-up] and third [at one month follow-up] auctions) and training conditions (Go, NoGo, Spaced, or Massed) on WTP within each pair type separately (Both Go, Both NoGo, Both Spaced or Both Massed). We conducted four separate 2 × 3 interactions rather than the full 2 × 2 × 3 interaction because items were selected to be in the different pairs based on WTP and thus initial differences in WTP between Go and NoGo or Spaced and Massed items are artificial and due to the item selection and pairing procedure rather than due to real differences. Running the regressions only on items that were paired together during the probe phase and by design matched for initial WTP is most appropriate. These interactions test whether the change in WTP over time is different for Spaced and Massed or Go and NoGo items. P values for the effects in the mixed models were calculated using the Kenward-Roger approximation for degrees of freedom [26].

## Results and Discussion

### Study 1

We conducted this study to test whether spacing CAT trials over two days while expanding the within-session lag from day 1 to day 2 and presenting Massed items only on day 2 will better preserve the choice of Go over NoGo items one week and one month after the end of training. We also tested whether this spacing and massing schedule induced a choice preference for Spaced over Massed items.

#### Go vs. NoGo

Massed Go items were chosen over Massed NoGo items at an immediate probe following cue-approach training (Both Massed black bar in Fig 4A, see Table 3 for all statistics). This is a replication of previous results demonstrating a shift in choice preferences in favor of Go items immediately following cue-approach training [3]. The preference for Massed Go over Massed NoGo items appeared to decrease over time, while choice of Massed Go was significant at the immediate probe, it decreased to non-significant at the one-month follow-up probe (Both Massed in Fig 4A). Similarly, Spaced Go items were chosen over Spaced NoGo items at an immediate probe following cue-approach training (Both Spaced black bar in Fig 4A, see Table 3 for all statistics). This preference for Spaced Go over Spaced NoGo items decreased but remained significant one month after the end of cue-approach training (Both Spaced white bar in Fig 4A). However, contrary to our prediction, there was no interaction between pair type (Both Spaced / Both Massed) and probe time (immediate/one-month follow-up) and no main effect of pair type on choices of Go items at probe. Taken together, these results suggest that employing the spacing schedule in study 1 was of only marginal benefit to help maintain choice of Go over NoGo items over the long term.

**Table 3:** Descriptive statistics for probe phase behavior in Study 1. Proportion choice of Go over NoGo (top) or choice of Spaced over Massed (bottom). Odds ratio (O.R) for choice of Go to NoGo (top) or Spaced to Massed (bottom). Confidence interval (C.I) on odds ratio and p-value for odds of choosing Go (top) or Spaced (bottom) item against equal odds. Interaction p-value (Xp) of pair type by probe time on odds of choosing Go to NoGo (top) or Spaced to Massed (bottom). Main effect p-value (ME p) of Spaced greater than Massed on choices of Go (top) or Go greater than NoGo on choices of Spaced (bottom).

| Pair Type | Time | P(Chose Go) | O.R | C.I | p-value | X p | ME p |
| --- | --- | --- | --- | --- | --- | --- | --- |
| Both | immediate | 70% | 3.23 | [1.71 6.09] | 0.0003 | 0.9 | 0.3 |
| Massed | 1 month | 58% | 1.45 | [0.93 2.26] | 0.1 |  |  |
| Both | immediate | 74% | 5.04 | [2.38 10.7] | < 0.0001 |  |  |
| Spaced | 1 month | 62% | 2.27 | [1.18 4.36] | 0.01 |  |  |

Table 3: Descriptive statistics for probe phase behavior in Study 1.
| Pair Type | Time | P(Chose Spaced) | O.R | C.I | p-value | X p | ME p |
| --- | --- | --- | --- | --- | --- | --- | --- |
| Both | immediate | 48% | 0.90 | [0.61 1.34] | 0.6 | 0.09 | 0.09 |
| NoGo | 1 month | 48% | 0.92 | [0.66 1.27] | 0.6 |  |  |
| Both Go | immediate | 53% | 1.13 | [0.78 1.65] | 0.5 |  |  |
|  | 1 month | 60% | 1.6 | [1.14 2.24] | 0.007 |  |  |

#### Spaced vs. Massed

There was no effect of spacing on choice of Spaced NoGo over Massed NoGo at the immediate probe. The lack of choice of Spaced NoGo over Massed NoGo was maintained at the one-month follow-up (Both NoGo in Fig 4B). Participants did not choose Spaced Go over Massed Go items at an immediate probe (Both Go in Fig 4B). However, the choice of Spaced Go over Massed Go increased at the one-month follow-up probe (rightmost white bar in Fig 4B). Weakly consistent with our predictions, there was a marginal interaction between pair type (Both Go / Both NoGo) and probe time (immediate/one-month follow-up) on choices of Spaced items at probe (Fig 4B, p = 0.09). There was also a marginal main effect of pair type (Go greater than NoGo) on choices of Spaced items. These results suggest that spacing cue-approach trials over two days and expanding the within-session lag from day 1 to day 2 induces a shift in preferences for Spaced over Massed items, but only if said items are associated with a Go tone during training.

#### Auction

There were no significant effects of spacing cue-approach training on the subjective value placed on food items. For all pair types of interest, item WTPs decreased equivalently over time (main effect of probe number [initial, one-week and one-month follow-ups] p’s < 0.0001, but no main effect of or interaction with item type [all p’s > 0.07] on WTP). The overall decrease in WTP over time is likely due to regression toward the mean, given that all items in this analysis were high value items based on the first auction. Regression to the mean in WTP measures over time has been previously reported [3]. These results suggest that spacing cue-approach training trials does not influence the subjective value placed on food items.

### Study 2

Because memory for Spaced items in study 1 could have been stronger due to primacy effects [27] and could have influenced choices [28], we conducted study 2 to test whether spacing CAT trials over two days while contracting the within-session lag from day 1 to day 2 and presenting Massed items only on day 1 (thus eliminating the primacy of Spaced item presentation) will help preserve the choice of Go over NoGo items one week and one month after the end of training. We also tested whether this spacing and massing schedule induced a choice preference for Spaced over Massed items.

#### Go vs. NoGo

Participants in study 2 chose Massed Go over Massed NoGo at an immediate probe, replicating previous findings. Choices of Massed Go items decreased significantly over time (Both Massed in Fig 5A, Table 4 for all statistics). Similarly, participants chose Spaced Go over Spaced NoGo items at a probe that took place immediately following cue-approach training (black bar Both Spaced in Fig 5A, see Table 4 for all statistics). This preference for Spaced Go over Spaced NoGo items decreased significantly one month after the end of CAT (white bar Both Spaced in Fig 5A). Again, contrary to our prediction, there was no interaction between pair type (Both Spaced / Both Massed) and probe time (immediate/one-month follow-up) on choices of Go items at probe. These results confirm findings in study 1 and suggest that spacing cue-approach training trials over two days and massing items on a single day does not significantly benefit maintenance of Go over NoGo item choice in the long term.

**Fig 5:**
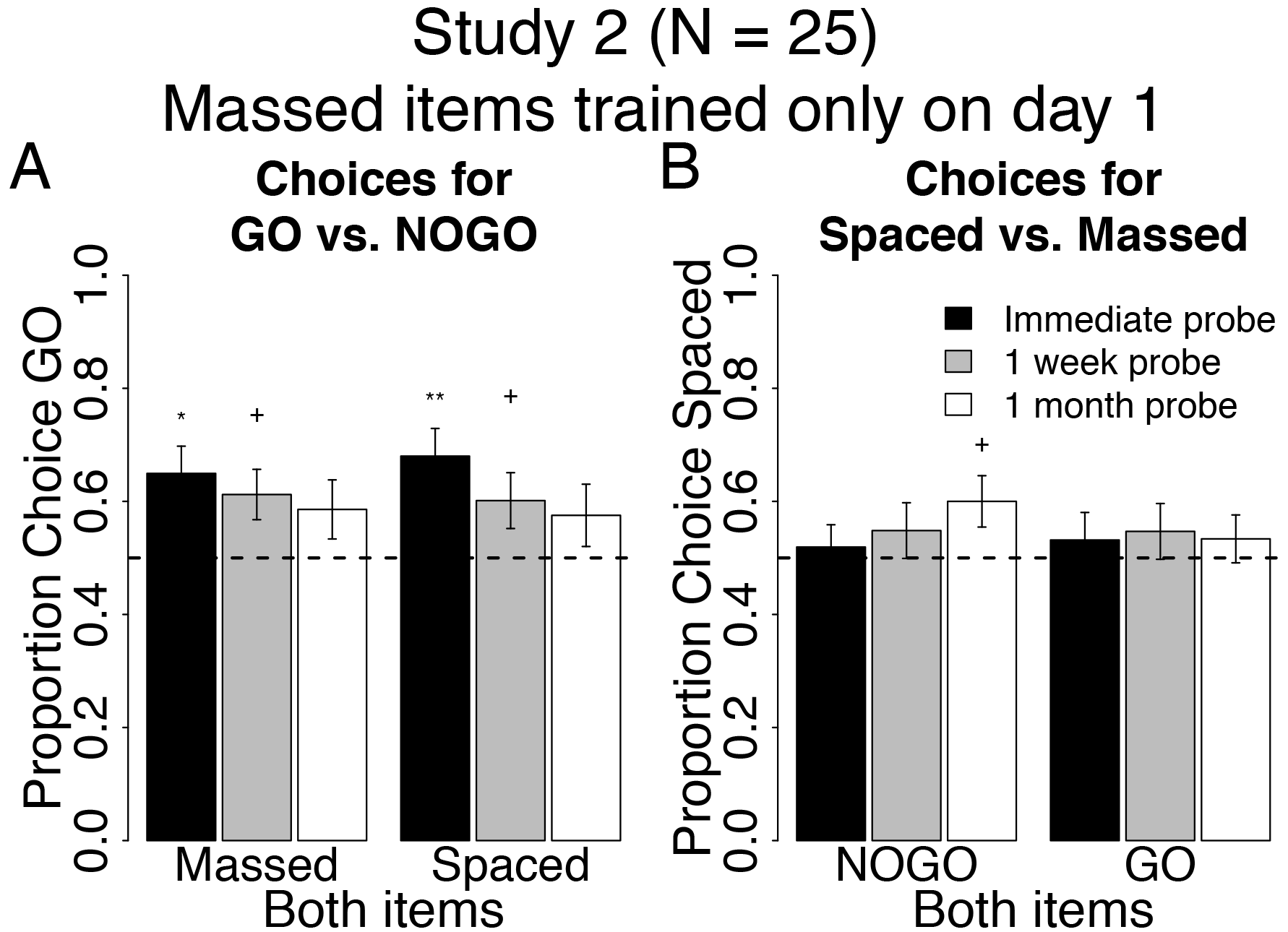
Behavioral results at probe for spacing of cue-approach training Study 2. Proportion choice of the Go vs. NoGo item (A) and proportion choice of the Spaced vs. Massed item (B) at probe immediately after training (black bars), one week (grey bars) and one month later (white bars). All error bars reflect one standard error of the mean (SEM). +: *p* < 0.05, *:*p* <;0.01, **: *p* < 0.001 in two-tailed repeated measures logistic regression for odds of choosing Go to NoGo (A) or Spaced to Massed (B) against equal odds.

**Table 4:** Descriptive statistics for probe phase behavior in Study 2. Proportion choice of Go over NoGo (top) or choice of Spaced over Massed (bottom). Odds ratio (O.R) for choice of Go to NoGo (top) or Spaced to Massed (bottom). Confidence interval (C.I) on odds ratio and p-value for odds of choosing Go (top) or Spaced (bottom) item against equal odds. Interaction p-value (X p) of pair type by probe time on odds of choosing Go to NoGo (top) or Spaced to Massed (bottom). Main effect p-value (ME p) of Spaced greater than Massed on choices of Go (top) or Go greater than NoGo on choices of Spaced (bottom).

| Pair Type | Time | P(Chose Go) | O.R | C.I | p-value | X p | ME p |
| --- | --- | --- | --- | --- | --- | --- | --- |
| Both | immediate | 65% | 2.23 | [1.32 3.77] | 0.003 | 0.3 | 0.9 |
| Massed | 1 month | 59% | 1.68 | [0.95 2.99] | 0.08 |  |  |
| Both | immediate | 68% | 2.62 | [1.53 4.49] | 0.0005 |  |  |
| Spaced | 1 month | 58% | 1.50 | [0.81 2.78] | 0.2 |  |  |

Table 4: Descriptive statistics for probe phase behavior in Study 2.
| Pair Type | Time | P(Chose Spaced) | O.R | C.I | p-value | X p | ME p |
| --- | --- | --- | --- | --- | --- | --- | --- |
| Both | immediate | 52% | 1.08 | [0.78 1.51] | 0.6 | 0.08 | 0.8 |
| NoGo | 1 month | 60% | 1.60 | [1.06 2.41] | 0.03 |  |  |
| Both | immediate | 53% | 1.17 | [0.75 1.84] | 0.5 |  |  |
| Go | 1 month | 53% | 1.18 | [0.81 1.71] | 0.4 |  |  |

#### Spaced vs. Massed

Participants chose Spaced NoGo and Massed NoGo items equivalently at an immediate probe (Both NoGo in Fig 5B). However, at the one-month follow-up probe, choice of Spaced NoGo over Massed NoGo items increased (white bar on the left in Fig 5B). There was no effect of spacing on choice of Spaced Go over Massed Go items at either the immediate or the one-month follow-up probes (Both Go in Fig 5B). There was a marginal interaction between pair type (Both Go / Both NoGo) and probe time (immediate/one-month follow-up) on choices of Spaced items at probe (p = 0.08), but this interaction was driven by a positive effect of time on choices of Spaced for Both NoGo items, rather than for Both Go items as predicted. There was also a marginal interaction between study (1 or 2) and probe time (immediate/one-month follow-up) on choices of Spaced items within Both NoGo trials (Both NoGo in Figs 4B vs. 5B, p = 0.07) and within Both Go trials (Both Go in Figs 4B vs. 5B, p = 0.09). These results suggest that spacing cue-approach trials over two days and reducing the within-session lag from day 1 to day 2 induces a shift in preferences for Spaced over Massed items, but only if said items are not associated with a Go tone during training.

#### Auction

Spacing CAT had no significant effect on the subjective value placed on food items. For all pair types of interest, item WTPs decreased equivalently over time (main effect of probe number [initial, one week and one-month follow-ups] p’s < 0.0001, but no main effect of or interaction with item type [all p’s >;0.4] on WTP). These findings confirm that spacing cue-approach training trials does not seem to influence the subjective value placed on food items.

#### Interim discussion and conclusions for Studies 1 & 2

In these studies, we found that spacing CAT trials did not have a lasting effect on choices of Go over NoGo, but did seem to induce a preference for Spaced over Massed items. There are several considerations in the design of studies 1 and 2 we would like to highlight: first, the comparisons between Spaced and Massed in these studies are largely testing lag effects rather than pure spacing effects. Lag effects tend to be weaker than spacing effects [29], perhaps masking more robust findings in studies 1 & 2. Second, a negative effect on spacing for longer compared to medium spacing or lag intervals has been previously found [29,30], which has been theorized to be due to failures to recognize repetitions as such [31]. Thus, the potential negative effect of longer lags on maintenance of choice preference shift in studies 1 & 2 may have in part driven the choice of Spaced over Massed items, given that Massed items have longer lags on the days when they were trained along with Spaced items. Furthermore, Spaced items appeared alone (no Massed items) on one of the days of training in both studies 1 & 2, potentially reducing the effectiveness of the spacing manipulation due to pure list effects. Encoding in pure lists has been shown to reduce or abolish spacing effects in free recall [32]. Finally, there was a difference in retention interval between Spaced and Massed items in Studies 1 & 2, which might also be tied to the primacy (study 2) or recency (study 1) of Spaced items.

Study 2 was conducted in order to test whether recency of Spaced items had an effect on choices of Spaced vs. Massed items. In Study 1, choices of Spaced Go over Massed Go items (white bar Both Go in Fig 4B) could be interpreted as a primacy rather than a spacing effect, given that Spaced items were seen earlier in the experiment on day 1 compared to Massed items, which were not seen until day 2 in Study 1. We found that choices of Spaced over Massed items increased as time elapsed in both studies, although this increase in Spaced choice over time effect shifted from Go items in Study 1 (primacy of Spaced items) to NoGo items in Study 2 (recency of Spaced items). These results highlight the potential importance of spacing cue-approach trials on long-term preference change. However, studies 1 and 2 might have suffered from several potential confounds discussed above and did not definitively demonstrate the hypothesized maintenance of basic Go over NoGo item choice effect over time. This could be due to the asymmetric training length between day 1 and day 2, different lags, and pure list effects. Therefore, we set out to address these confounds and further test the effects of spacing on the persistence of Go over NoGo choice preference by equating training length on day 1 and day 2, employed zero-lag massing schedule to
test pure spacing rather than lag effects, and distributed Massed item training over both days to address pure list effects in Study 3.

### Study 3

In studies 1 and 2, massed items had a non-zero lag, i.e. had a number of intervening other items in between presentations of the same massed item. Zero-lag massing is defined as training repetitions with zero within-session lag, meaning repetitions for a particular Massed item during training are consecutive, with no other item presentations in between. In studies 1 and 2, Massed items had half the within-session lag of Spaced items (which was higher than zero), but all training of Massed items occurred on only one of the two training days. We conducted study 3 to test whether spacing cue-approach trials over two days while maintaining the same within-session lag from day 1 to day 2, zero-lag massing items (training with zero within-session lag) and presenting Massed items on both day 1 and day 2 helped preserve the choice of Go over NoGo items one week and one month after the end of training. We also tested whether this spacing and massing schedule induced a choice preference for Spaced over Massed items.

#### Go vs. NoGo

Participants in study 3 chose Spaced Go over Spaced NoGo items consistently over time (Both Spaced in Fig 6A, see Table 5 for all statistics). However, participants consistently had no preference for Massed Go over Massed NoGo items over time (Both Massed in Fig 6A), which is inconsistent with previous findings. Contrary to our prediction, there was no interaction between pair type (Both Spaced / Both Massed) and probe time (immediate / one-month follow-up) on choices of Go over NoGo items. However, consistent with our secondary prediction, we found a main effect of pair type (Both Spaced greater than Both Massed, p = 0.03), but no main effect of probe time (immediate / one-month follow-up) on choices of Go over NoGo items.

**Fig 6:**
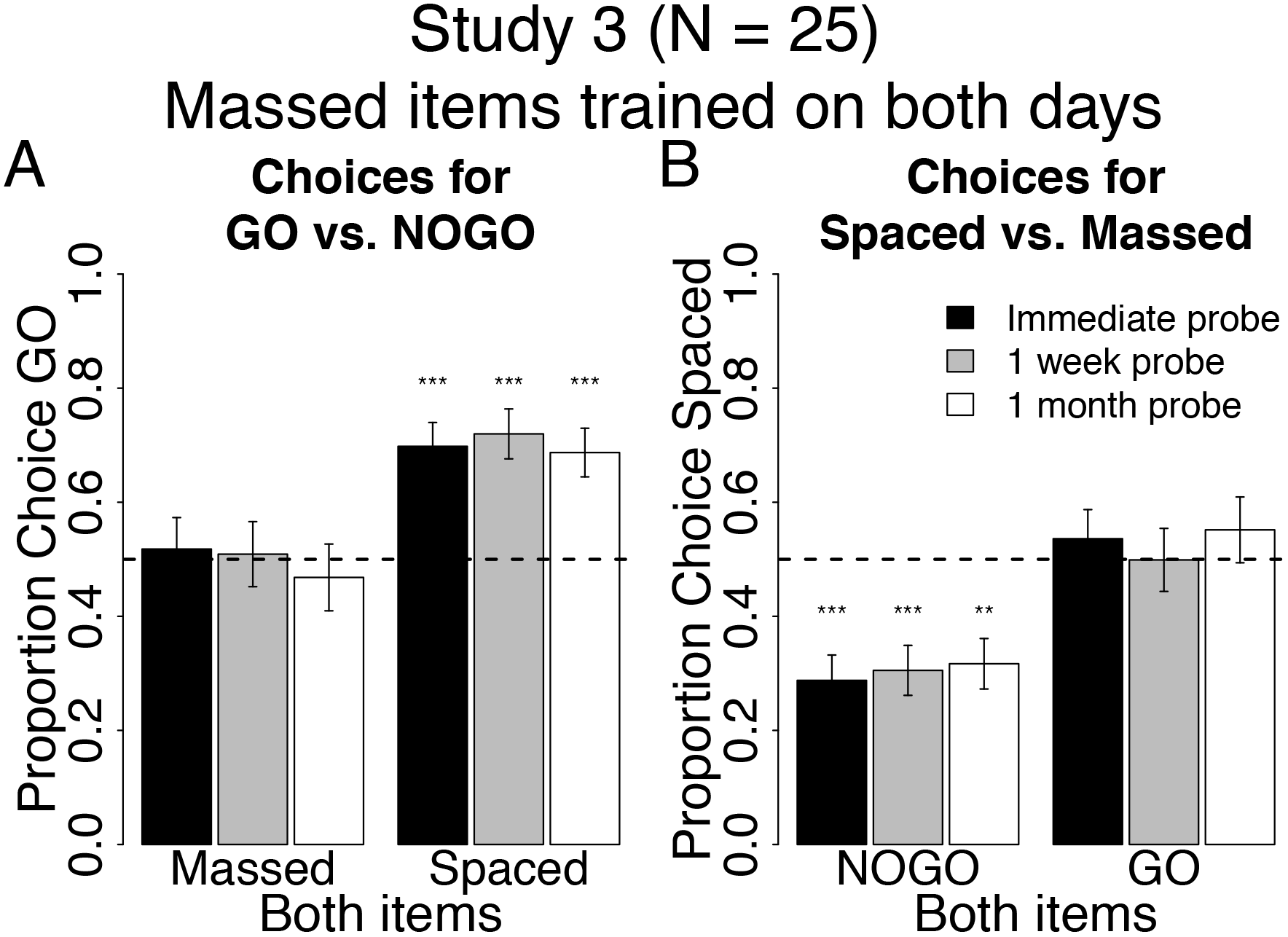
Behavioral results at probe for spacing of cue-approach training Study 3. Proportion choice of the Go vs. NoGo item (A) and proportion choice of the Spaced vs. Massed item (B) at probe immediately after training (black bars), one week (grey bars) and one month later (white bars). All error bars reflect one standard error of the mean (SEM). **: *p* < 0.001, ***: *p* < 0.0001 in two-tailed repeated measures logistic regression for odds of choosing Go to NoGo (A) or Spaced to Massed (B) against equal odds.

**Table 5:** Descriptive statistics for probe phase behavior in Study 3. Proportion choice of Go over NoGo (top) or choice of Spaced over Massed (bottom). Odds ratio (O.R) for choice of Go to NoGo (top) or Spaced to Massed (bottom). Confidence interval (C.I) on odds ratio and p-value for odds of choosing Go (top) or Spaced (bottom) item against equal odds. Interaction p-value (X p) of pair type by probe time on odds of choosing Go to NoGo (top) or Spaced to Massed (bottom). Main effect p-value (ME p) of Spaced greater than Massed on choices of Go (top) or Go greater than NoGo on choices of Spaced (bottom).

| Pair Type | Time | P(Chose Go) | O.R | C.I | p-value | X p | ME p |
| --- | --- | --- | --- | --- | --- | --- | --- |
| Both | immediate | 52% | 1.14 | [0.64 2.04] | 0.7 | 0.4 | < 0.0001 |
| Massed | 1 month | 47% | 0.85 | [0.46 1.57] | 0.6 |  |  |
| Both | immediate | 70% | 2.77 | [1.76 4.38] | < 0.0001 |  |  |
| Spaced | 1 month | 69% | 2.64 | [1.66 4.19] | < 0.0001 |  |  |

Table 5: Descriptive statistics for probe phase behavior in Study 3.
| Pair Type | Time | P(Chose Spaced) | O.R | C.I | p-value | X p | ME p |
| --- | --- | --- | --- | --- | --- | --- | --- |
| Both | immediate | 29% | 0.32 | [0.19 0.53] | < 0.0001 | 0.7 | 0.005 |
| Massed | 1 month | 32% | 0.38 | [0.23 0.62] | 0.0001 |  |  |
| Both | immediate | 54% | 1.20 | [0.72 1.99] | 0.5 |  |  |
| Spaced | 1 month | 55% | 1.24 | [0.67 2.30] | 0.5 |  |  |

#### Spaced vs. Massed

Participants consistently chose Massed NoGo over Spaced NoGo items at all probes (Both NoGo in Fig 6B). There was no effect of spacing on choice of Spaced Go over Massed Go items at any probe (Both Go in Fig 6B). Again, contrary to our prediction, there was no interaction between pair type (Both Go / Both NoGo) and probe time (immediate / one-month followup) on choices for Spaced over massed. There was a main effect of Both Go greater than Both NoGo on choices of Spaced, but this was driven by a bias to choosing Massed NoGo items, which is not what we had predicted.

#### Auction

There were no significant effects of spacing cue-approach training on the subjective value placed on food items. For all pair types of interest, item WTPs decreased equivalently over time (main effect of probe number [initial, one week and one-month follow-ups] p’s < 0.0001, but no main effect of or interaction with item type [all p’s > 0.1] on WTP). These results replicate findings from studies 1 and 2, which confirm that spacing cue-approach training trials had no effect on the subjective value placed on food items.

### Study 4

Study 4 is a direct, pre-registered replication of study 3 [33]. The registration can be found on the open science framework at https://osf.io/pgyrv/.

#### Go vs. NoGo

The pattern of behavior for choices of Go vs. NoGo items in study 4 is very similar to that in study 3. Participants chose Spaced Go over Spaced NoGo items consistently over time (Both Spaced in Fig 7A, see Table 6 for all statistics). However, participants consistently had no preference for Massed Go over Massed NoGo items over time (Both Massed in Fig 7A). Again, contrary to our prediction, but consistent with study 3, there was no interaction between pair type (Both Spaced / Both Massed) and probe time (immediate / one-month follow-up) on choices of Go over NoGo items. However, consistent with our secondary prediction and study 3, we found a main effect of pair type (Both Spaced greater than Both Massed, p = 0.001) on choices of Go over NoGo items.

**Table 6:** Descriptive statistics for probe phase behavior in Study 4. Proportion choice of Go over NoGo (top) or choice of Spaced over Massed (bottom). Odds ratio (O.R) for choice of Go to NoGo (top) or Spaced to Massed (bottom). Confidence interval (C.I) on odds ratio and p-value for odds of choosing Go (top) or Spaced (bottom) item against equal odds. Interaction p-value (X p) of pair type by probe time on odds of choosing Go to NoGo (top) or Spaced to Massed (bottom). Main effect p-value (ME p) of Spaced greater than Massed on choices of Go (top) or Go greater than NoGo on choices of Spaced (bottom).

| Pair Type | Time | P(Chose Go) | O.R | C.I | p-value | X p | ME p |
| --- | --- | --- | --- | --- | --- | --- | --- |
| Both | immediate | 56% | 1.30 | [0.97 1.74] | 0.08 | 0.1 | 0.001 |
| Massed | 1 month | 51% | 1.09 | [0.79 1.49] | 0.6 |  |  |
| Both | immediate | 73% | 3.05 | [2.20 4.21] | < 0.0001 |  |  |
| Spaced | 1 month | 64% | 2.29 | [1.37 3.84] | 0.001 |  |  |

Table 6: Descriptive statistics for probe phase behavior in Study 4.
| Pair Type | Time | P(Chose Spaced) | O.R | C.I | p-value | X p | ME p |
| --- | --- | --- | --- | --- | --- | --- | --- |
| Both | immediate | 44% | 0.73 | [0.48 1.13] | 0.2 | 0.01 | 0.01 |
| Massed | 1 month | 52% | 1.07 | [0.76 1.52] | 0.7 |  |  |
| Both | immediate | 61% | 1.69 | [1.24 2.31] | 0.0009 |  |  |
| Spaced | 1 month | 60% | 1.73 | [1.11 2.70] | 0.02 |  |  |

#### Spaced vs. Massed

Participants in study 4 consistently chose Spaced Go over Massed Go items at all probes (Both Go in Fig 7B). There was no effect of spacing on choice of Spaced NoGo over Massed NoGo items at any probe (Both NoGo in Fig 7B). There was a significant interaction between pair type (Both Go / Both NoGo) and probe time (immediate / one-month follow-up) on choices for Spaced over massed, but the interaction was driven by an increasing rate of choice for Spaced NoGo and a decreasing rate of choice for Spaced Go, which is different from what we predicted. However, consistent with our secondary prediction, there was a main effect of Both Go greater than Both NoGo on choices of Spaced Go items.

#### Auction

There were no significant effects of spacing cue-approach training on the subjective value placed on food items. For all pair types of interest, item WTPs decreased equivalently over time (main effect of probe number [initial, one week and one-month follow-ups] p’s < 0.001, but no main effect of or interaction with item type [all p’s > 0.1] on WTP). These results replicate findings from studies 1, 2 & 3, which confirm that spacing cue-approach training trials had no effect on the subjective value placed on food items.

#### Interim discussion and conclusions for Studies 3 & 4

In studies 3 & 4, we addressed the confounds present in studies 1 & 2 and tested the efficacy of spacing CAT trials in helping maintain the preference for Go over NoGo foods as well as preference for Spaced over Massed foods. Results suggest that spacing cue-approach trials over two days and maintaining the same within-session lag across days offer significant benefit to the maintenance of Spaced Go over Spaced NoGo choice over time (Both Spaced in Figs 6A and 7A). Zero-lag massing cue-approach training trials - i.e. presenting Massed items consecutively with no intervening items - eliminates the Go choice effect altogether (Both Massed in Figs 6A and 7A). Zero-lag massing items seems to induce a strong and lasting preference for Massed over Spaced items, but only if said items were not associated with a Go cue during cue-approach training (Both NoGo versus Both Go in Fig 6B). However, this preference for Massed NoGo over Spaced NoGo is not robust and did not replicate in study 4 (Both NoGo in Fig 7B). The lack of an expected Massed Go over Massed NoGo choice preference (Both Massed in Fig 6B) in study 3 could be due to an increase in preference for Massed NoGo items (as seen for Both Massed in Fig 6B) that counteracts the regularly induced preference for Go items (Figs 4A and 5A). However, the lack of preference for Massed Go over Massed NoGo (Both Massed in Fig 7A) replicated in study 4, despite the lack of preference for Massed NoGo over Spaced NoGo (Both NoGo in Fig 6B) to counteract it as was the case in Study 3.

**Fig 7:**
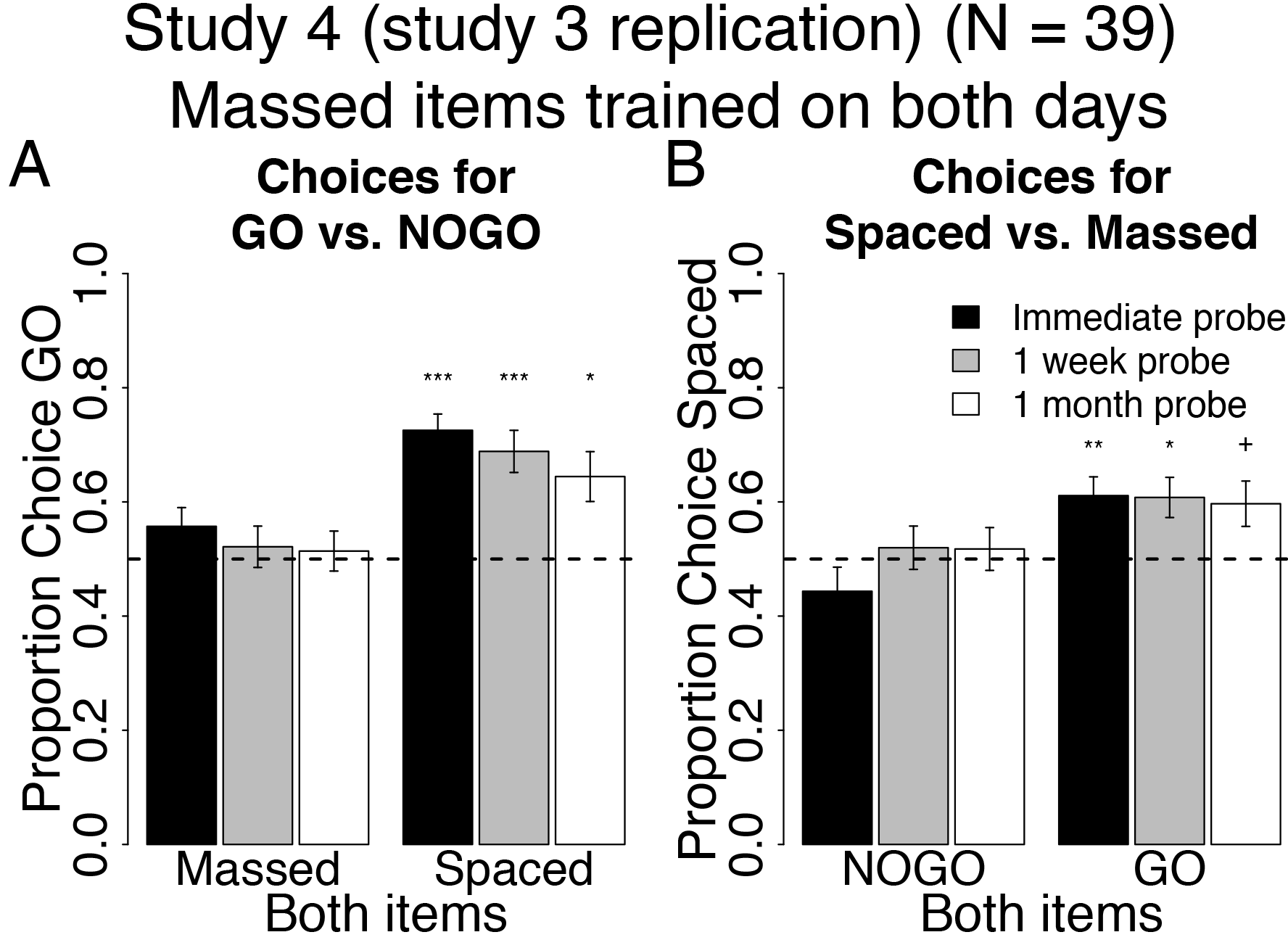
Behavioral results at probe for spacing of cue-approach training Study 4 (direct replication of study 3). Proportion choice of the Go vs. NoGo item (A) and proportion choice of the Spaced vs. Massed item (B) at probe immediately after training (black bars), one week (grey bars) and one month later (white bars). All error bars reflect one standard error of the mean (SEM). +: *p* < 0.05, *:*p* < 0.01, **: *p* < 0.001, ***: *p* < 0.0001 in two-tailed repeated measures logistic regression for odds of choosing Go to NoGo (A) or Spaced to Massed (B) against equal odds.

Studies 3 & 4 employ zero-lag massing, where all trials for a particular Massed Go item are consecutive with no intervening trials involving other trial types or items. In these studies, after two or three of the 12 presentations of a particular Massed Go item, participants learn that the next several trials will also be Go trials involving the same item. Thus, Massed Go trials quickly lose their novelty and surprise in Studies 3 & 4 compared to the designs in studies 1 and 2 or in the previous standard version of CAT in Schonberg et al. [3]. The potential lack of sustained attention toward Massed Go items might contribute to a lack of previously seen and expected preference shift for Massed Go over Massed NoGo in Study 3 and is consistent with findings from our group showing that presenting Go items in blocks of Go trials eliminates the expected preference for Go over NoGo items [4].

#### General discussion

Previous work has established CAT as a reliable method to influence choice by targeting automatic processes rather than relying on effortful control of behavior [1,3]. The shift in preference for appetitive snack food items was maintained over two months following the longest training period and for one month in other samples [3]. In the current studies, we sought to improve the long-term maintenance of a change in choice behavior by spacing CAT trials over two consecutive days. The distributed practice effect is one of the most robust findings in the memory literature [18,21,34]. Spacing strategies have been applied during extinction with the goal of preventing the return of fear in the long-term [22,35], but these strategies have not been widely adopted in the appetitive domain. Here, we apply the principles of distributed practice and space CAT trials over two days in three studies to test the effectiveness of spacing strategies on the maintenance or change over time in the shift in choice preferences. The three studies presented here reveal a benefit of spacing CAT trials over two days to the long-term shift in choice preferences and the persistence of the standard Go choice effect over time, depending on the spacing and massing schedule.

#### Studies 1 & 2

In studies 1 and 2, Massed items appeared only on one day (only on day 2 in Study 1 and only on day 1 in Study 2). This massing schedule resulted in an increase in choice preference for Spaced items over time. These results suggest that spacing presentations of some items over two days while massing presentations of other items on a single day (resulting in a longer training session on one day than the other) modulated later preference for Spaced over Massed items one month after the end of training. The order in which Massed items, or the expanding or contracting within-session lag for Spaced items, seems to have an impact on choice of Spaced over Massed items that interacts with their Go/NoGo status. Expanding the within-session lag from day 1 to day 2 as in Study 1 (average lag for Spaced items on day 1 = 24 and on day 2 = 72) increases the choice of Spaced Go over Massed Go, but does not benefit the choice of Spaced NoGo over Massed NoGo items over time. On the other hand, contracting the within-session lag for Spaced items from day 1 to day 2 as in Study 2 (average lag for Spaced items on day 1 = 72 and on day 2 = 24) boosts the choice of Spaced NoGo over Massed NoGo, but does not increase the choice of Spaced Go over Massed Go items over time.

Massing cue-approach training trials on a single day and keeping training length unbalanced across the two days seems to not serve the maintenance of the Go choice effect. There was no significant choice of Massed Go over Massed NoGo at the one-month follow-up. There was also only weakly significant choice of Spaced Go over Spaced NoGo at the one-month follow-up in Study 1. The interpretation of results from studies 1 & 2 are complicated by a number of confounds such as testing lag effects that are weaker than pure spacing effect, differing lag intervals which may artificially enhance or impede detection of spacing effects, and pure list effects which tend to weaken spacing effects. These confounds are discussed in the interim discussion for studies 1 & 2 above.

#### Studies 3 & 4

In studies 3 & 4, half the Massed items appeared on each of the two training days, addressing the pure list confound, and all the presentations of a particular Massed item were presented on one day, consecutively, with no intervening other items in between presentations (zero-lag massing), allowing us to test for pure spacing effects rather than lag effects. Spaced item presentations were spread over the two days as in Study 1 and 2, resulting in the same training session length on day 1 and day 2. This massing schedule revealed a robust maintenance of the standard Go choice effect over time. These results suggest that spacing cue-approach training over two days using this schedule benefits the preservation of a shift in choice preferences in favor of Spaced Go items over one month following CAT.

Zero-lag massing training trials so that trials are presented consecutively, with no intervening other items (i.e. zero within-session lag), appears to have eliminated the Go choice effect. Zero-lag massing trial presentations produces a lasting preference for Massed NoGo items, with consistent choice of Massed NoGo over Spaced NoGo at all three probes, but this effect was not replicated in Study 4. This preference for zero-within-session-lag Massed NoGo items in Study 3 may be interfering with the regularly observed preference for Go over NoGo items and may explain the lack of choice of Massed Go over Massed NoGo at all probes in Study 3. But given that the lack of preference for Massed Go over Massed NoGo replicated in Study 4, it seems more likely that predictability of the Go cue among a block of consecutive Massed Go trials is the cause for the lack of Go preference when both items are Massed, which is consistent with previous findings when we trained Go in predictable blocks [4]. Notice that there is no preference for Massed NoGo over Spaced NoGo items in Study 4. Studies 3 & 4 are inconsistent as to the benefit of zero-lag massed training trials with no lag for Massed items and maintaining the same within-session lag for Spaced items on both training days in (with an average lag of 48 on both day 1 and day 2) on the choice of Spaced Go over Massed Go items. Both studies show a main effect for Both Go greater than Both NoGo for choice of Spaced over Massed, but the effect is driven as predicted by higher choices for Spaced Go only in Study 4.

#### Spacing effects on stimulus value

Although spacing CAT trials can influence choice, it does not seem to have an effect on the subjective value placed on food items. We have previously suggested that attentional and memory mechanisms are at play during CAT [3–5], although the contribution of each to choice and valuation remain unknown. Further work is needed to fully understand the role of memory during this task and its influence on choice behavior and valuation. Better characterization of the role of memory in this task may help optimize the spacing schedule during cue-approach training to yield the longest maintenance duration of behavioral change.

#### Further considerations

The cue-approach task is related to extensive work on the attentional boost effect, which shows a later memory advantage for task-irrelevant information when presented concurrently with a behaviorally relevant stimulus [for review, see 13]. However, to our knowledge, no attentional boost study has shown a long-term memory advantage over weeks or months following the distributed practice of attentional boost training.

Previous research has determined that a lag of one day between study sessions was optimal for retention intervals of about a week to one month for verbal learning tasks [18]. For practical reasons, we chose to bring participants back for the final probe four weeks after the end of cue-approach training, thus opting for a between-session lag of one day in the current studies. This conforms to conventional wisdom in the field that optimal between-study episode lag is around 10-20% of the retention interval [17]. Further studies that vary between-session lag for Spaced items will be needed to define the boundaries of the spacing effect in the cue-approach task.

Most studies on the spacing effect to date employ motor skill or verbal learning tasks and to our knowledge none have explored the spacing effect using a non-reinforced associative task such as the cue-approach task employed here. Pashler et al. [36] have reported a lack of spacing effect using perceptual categorization tasks. Although more research is needed on the topic, it seems not all forms of learning benefit from spacing. Given the associative nature of the task employed here, we believe that spacing principles are likely to be applicable to the cue-approach task. More studies employing spacing in other forms of non-reinforced learning tasks are needed to fully understand the task characteristics that make spacing effective in improving long term performance in these types of tasks.

### Conclusions

In conclusion, we propose that spacing CAT trials may help maintain the change in preference for appetitive food over the long term and that different spacing schedules can be employed to optimize the desired long-term changes. Although not widely adopted by clinicians, spacing strategies have proven useful in the treatment of anxiety disorders [22]. Several researchers have for some time advocated the implementation of spacing strategies in instruction, given its clear advantage for long-term memory retention and its applicability to academic goals [17,37]. Here we show that similar strategies may potentially be useful for attaining more common behavioral change goals such as maintaining healthy weight.

## Acknowledgements

This research was funded by National Institutes of Health grant R01AG041653 awarded to RAP and by Israeli Science Foundation (ISF) grant 1798/15 to TS. The authors would like to thank Jave Del Rosario and Kelly Jameson for help with data collection.

